# Bilayer tension-induced clustering of the UPR sensor IRE1

**DOI:** 10.1101/2023.05.26.542528

**Authors:** Md Zobayer Hossain, Wylie Stroberg

## Abstract

The endoplasmic reticulum acts as a protein quality control center where a range of chaperones and foldases facilitates protein folding. IRE1 is a sensory trans-membrane protein that transduces signals of proteotoxic stress by forming clusters and activating a cellular program called the unfolded protein response (UPR). Recently, membrane thickness variation due to membrane compositional changes have been shown to drive IRE1 cluster formation, activating the UPR even in the absence of proteotoxic stress. Here, we demonstrate a direct relationship between bilayer tension and UPR activation based on IRE1 dimer stability. The stability of the IRE1 dimer in a (50%DOPC-50%POPC) membrane at different applied bilayer tensions was analyzed via molecular dynamics simulations. The potential of mean force for IRE1 dimerization predicts a higher concentration of IRE1 dimers for both tensed and compressed ER membranes. This study shows that IRE1 may be a mechanosensitive membrane protein and establishes a direct biophysical relationship between bilayer tension and UPR activation.

**Highlights:**

- Mechanical perturbation of the ER membrane favor oligomerization.
- Both tension and compression promote IRE1 dimer formation.
- IRE1 is mechanosensitive, potentially the UPR to changes in ER membrane tension and compression.

## 1 Introduction

The endoplasmic reticulum (ER) performs a range of functions including protein folding, synthesis and transport, lipid metabolic regulation and calcium storage [Westrate et al., 2015, Clapham, 2007, Hebert et al., 2005, Fagone and Jackowski, 2009, Braakman and Hebert, 2013, Reid and Nicchitta, 2015, Rapoport, 2007]. Additionally, the ER acts as a protein quality control center where a range of chaperones and foldases facilitates protein folding. The scarcity of molecular chaperones relative to the folding demand of the nascent proteins in the ER creates a state where proteins fail to fold correctly. These misfolded/unfolded proteins accumulate in the ER lumen induces proteotoxic stress. In order to diminish proteotoxic stress and restore proteostasis in the ER, a broad set of responses called the unfolded protein response (UPR) is activated [Ron and Walter, 2007]. The UPR performs three major functions: suspension of protein translation, degradation of the misfolded protein, and increased folding capacity through the upregulation of genes encoding molecular chaperones. If the UPR fails to restore balance, apoptotic signals initiate cell death [Szegezdi et al., 2006, Hetz, 2012]. Proteotoxic stress and activation of the UPR are associated with many diseases including neurodegenerative diseases [Hetz and Saxena, 2017], type 2 diabetes [Decio L and Miriam, 2010, Allen and David, 2010], obesity [Gökhan S, 2010] and cancer [Xuemei et al., 2011].

Three proteins in the ER membrane transduce signals of ER stress. These are IRE1 (inositol requiring enzyme 1), PERK (double-stranded RNA-activated protein kinase [PKR]-like ER kinase), and ATF6 (Activating Transcription Factor 6) [Gardner et al., 2013]. Each transducer activates different UPR pathways to control proteostasis. In the case of IRE1, unfolded and misfolded protein in ER lumen facilitates IRE1 clustering by sequestering the chaperon BiP and stabilizing IRE1 oligomers [Calfon et al., 2002]. This clustered IRE1 produces transcription factors through splicing of mRNA HAC1/XBP1 to control the expression of UPR target genes [Yoshida et al., 2001, Cox and Peter, 1996]. The IRE1 branch of the UPR balances protein folding load by upregulating ER chaperons and decreasing translation of ER membrane-localized mRNA through Regulated IRE1-Dependent Decay (RIDD).

Recently, membrane aberrancies have been shown to induce IRE1 clustering and UPR activation in the absence of proteotoxic stress [Halbleib et al., 2017]. Membrane aberrancies due to compositional changes affects membrane thickness. So, membrane thickness mediated IRE1 clustering exhibits a mechanism of UPR activation in the absence of proteotoxic stress. Membrane thickness mediated interaction is also in regulation of several membrane protein’s function such as MscL gating [Perozo et al., 2002], gramicidin signalling [Elliott et al., 1983], pseudoequilibrium of Rhodospin [Brown, 1994] etc.

The difference between the hydrophobic section of the transmembrane domain of membrane proteins and the thickness of the bilayer is known as hydrophobic mismatch [Mouritsen and Bloom, 1984, Sperotto and Mouritsen, 1988, Sperotto et al., 1989, Killian and Nyholm, 2006]. Hydrophobic mismatch between transmembrane domains of protein and lipid results in local variations of membrane thickness depression around transmembrane proteins [Schmidt et al., 2008]. Altered membrane thickness lipid compositional changes leads to change in the hydrophobic mismatch of embedded proteins. The elliptical depression around IRE1 is due to an amphipathic portion of IRE1 that is constrained to sit at the interface between ER membrane and lumen. The amphipathic portion of the IRE1 depresses the membrane to create a favorable environment for clustering, activating UPR [Halbleib et al., 2017]. As the thickness of the bilayer increases upon compositional changes, the local deformation around the monomer must increase. The resulting increase in chemical potential energy for monomer can be ameliorated by forming dimers or higher order oligomers which allow the deformed regions around the proteins to overlap. Hence, membrane thickness due to compositional changes favor IRE1 clustering.

Similar to compositional changes, direct application of bilayer tension alters membrane thickness [Rawicz et al., 2000, Muddana et al., 2011, Klauda et al., 2010, Reddy et al., 2012]. Membrane tension is linked to several cell functions like endocytosis and exocytosis [Pontes et al., 2017]. In ER, membrane tension influences lipid budding and [Ben M’barek et al., 2017]. Cytoskeletal forces and lipid composition contribute to the tension in the membrane [Pontes et al., 2017]. Similarly, tension in ER will also be also determined by cytoskeletal forces and lipid composition. Tension applied to membrane surface changes area per lipid and membrane thickness, which impacts the overall packing density of lipids and proteins in the ER. As a result, ER tension will reorder the protein-lipid arrangement. Since bilayer tension is instrumental to changes in membrane thickness and packing density, We hypothesize that it also modulates the stability of IRE1 clusters.

This study determines the effect of membrane tension on IRE1 clustering through coarse-grained molecular dynamics simulations. These simulations demonstrate that IRE1, in addition to responding to protein concentration in the ER lume and lipid composition in the ER membrane, is also sensitive to mechanical perturbations to the ER membrane. The free energy difference between the IRE1 dimer and monomer states is used as an index for assessing the stability of IRE1 dimers under various tensions. In Section 2, we discuss molecular dynamics simulation techniques used to capture behaviour of the transmembrane portion of IRE1 in the ER. In Section 3, we present findings of how membrane tension alters stability of IRE1 dimers. In Section 4, we interpret the free energy landscape of the IRE1 dimerization in the tensed ER to suggest a biophysical relationship between tension and UPR. Finally, in Section 5, we conclude our study.

## 2 Methods

In the following section, simulation techniques and parameters used to calculate free energy landscapes of IRE1 dimerization in the ER are discussed in detail. First, we model the atomic structure of the transmembrane portion of IRE1, IRE1^516-571^. After validation of the atomistic structure, we built a coarse-grained structure of IRE1^516-571^ from the atomistic model. To estimate the free energy difference we then used replica exchange-umbrella sampling.

### 2.1 Atomistic modeling of IRE1 sensor peptide

The transmembrane portion of IRE1^516-571^ was adopted from the study of Halbleib et al. [Halbleib et al., 2017]. The 56 amino acid long sequence “516-SRELD EKNQNSLLLK FGSLVYRIIE TGVFLLLFLI FCAILQRFKI LPPLYVLLSK I-571” consists of a transmembrane helix and an amphipathic helix [Halbleib et al., 2017]. The IRE1 peptide was built from the sequence via the molefacture plugin of Visual Molecular Dynamics (VMD) [Humphrey et al., 1996]. IRE1^516-571^ was inserted into an equilibrated lipid bilayer of (50%-50%) DOPC-POPC via the membrane builder of the CHARMM-GUI [Jo et al., 2008, Brooks et al., 2009, Lee et al., 2016, Jo et al., 2009, Lee et al., 2019]. This membrane protein complex consists of a single IRE1^516-571^ monomer, 310 DOPC(1,2-Dioleoyl-sn-glycero-3-phosphocholine), 310 POPC (1-Palmitoyl-2-oleoyl-sn-glycero-3-phosphocholine) and 33702 TIP3P water molecules. For ionic balance, 91 sodium and 97 chloride ions were also inserted. The simulation box dimensions are 14.8 nm x 14.8 nm x 9.27 nm. The CHARMM36 force field was used for all simulations. After energy minimization, the model structure was first equilibrated while constraining the positions of lipid heads and protein. Another constraint was used to keep the water out of the hydrophobic core of the structure. After equilibration of 0.375 ns with time step 1 fs, the timestep was increased to 2 fs, and the system was equilibrated for an additional 0.5 ns. Next, a production run of 200 ns was carried out without constraints. All the simulations were run in the NPT ensemble by applying a Langevin thermostat at 303.15 K and Nose Hoover-Langevin piston barostat at 1 bar.

### 2.2 Coarse-grained IRE1 dimer

Large atom numbers (*≈* 180000) and time step of 2 fs makes atomistic simulations too expensive to achieve adequate sampling for replica exchange umbrella sampling simulations. On the other hand, reduced atom numbers and larger time steps make coarse-grained molecular dynamics simulation computationally significantly less expensive than the atomistic simulation. Hence, we opted for coarse-grained simulation for our study. The equilibrated structure of IRE1^516-571^ was extracted from the atomistic model. Two identical replicas of IRE1^516-571^ were placed such that the distance between carbon-*β* atoms of F544 residues of each monomer was 0.7 nm using VMD [Humphrey et al., 1996]. Monomers were oriented in an X-like configuration that has been shown to be the orientation of assembled full length IRE1 [Väth et al., 2021]. The dimer was coarse-grained using CHARMM GUI’s MARTINI Maker to produce a coarse-grained membrane protein complex [Hsu et al., 2017]. The coarse-grained model consists of 2 IRE1^516-571^ monomers, 574 DOPC and 574 POPC, and 22775 MARTINI water molecules. 270 negative ions and 270 positive ions were inserted into the system to make the system neutral. The dimension of the model is 20 nm x 20 nm x 10.15 nm. The MARTINI force field 2.0 was used for all interactions [Marrink et al., 2007]. After energy minimization, a short equilibration run was performed with the positions of lipid heads and proteins constrained. Next, the unconstrained system was equilibrated for 200 ns. All simulations were run in the NPT ensemble by applying a Langevin thermostat and Nose-Hoover barostat at 303.15 K and 1 bar.

### 2.3 Molecular dynamics and analysis softaware

The molecular simulation package NAMD was used for all molecular dynamics simulation [Phillips et al., 2020]. All visual representations were created via VMD [Humphrey et al., 1996]. For analysis of membrane thickness, the MEMBPLUGIN of VMD was used [Guixà-González et al., 2014]. The distance between coarse-grained lipid heads PO4 of the top and bottom leaflet of the membrane was defined as the membrane thickness. The colvar module was used to define the reaction coordinate and run steered molecular dynamics in NAMD [Fiorin et al., 2013]. Python 3.0 was used for data analysis and visualization. WHAM, an implementation of the weighted histogram analysis method, was used to calculate free energies from umbrella sampling simulations [Grossfield].

### 2.4 Replica-exchange umbrella sampling

Replica-exchange umbrella sampling (REUS) was implemented with the replica-exchange module of NAMD. The colvar module was used to specify the reaction coordinate and to record trajectory data. The root mean square of the inter-particle distance between backbone coarse-grained beads of IRE1 monomers, *d*_rms_, was used as the reaction coordinate. Even though *d*_rms_ is a 1D reaction coordinate, it was chosen specifically to capture some rotational degree of freedom along with the center of mass separation. *d*_rms_ is defined as

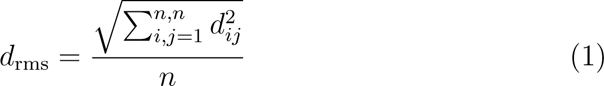

where *d_ij_*is the distance between backbone bead i of monomer A and bead j of monomer B and n is the number of backbone beads considered in each of the monomers.

From the simulation of IRE1^516-571^ monomer in the ER, we have found three distinct portion of IRE1^516-571^: IRE1^526-544^ sits on top of the lipid surface on the lumen side, IRE1^545-552^ is fully inserted into the lipids, and IRE1^553-571^ situated close to the cytosolic side (Fig. 1). To capture the movement with optimized computational expense, the backbone beads of IRE1^544-552^ were used in calculating *d*_rms_. Replica-exchange umbrella sampling was implemented with 35 evenly spaced bins over a range of 0.5*∼*7 nm (0.2 nm/bin). Replicas for the various bins were prepared using SMD with a spring constant of 25 kcal/mol^Å2^. In replica-exchange umbrella sampling, a harmonic restraint of 2.5 kcal/mol^Å2^ was applied to sample the replicas in various bins. Convergence analysis of replica-exchange umbrella sampling simulations are added in the Supplementary Material (Fig. S1,S2,S3,S4).

**Fig. 1:**
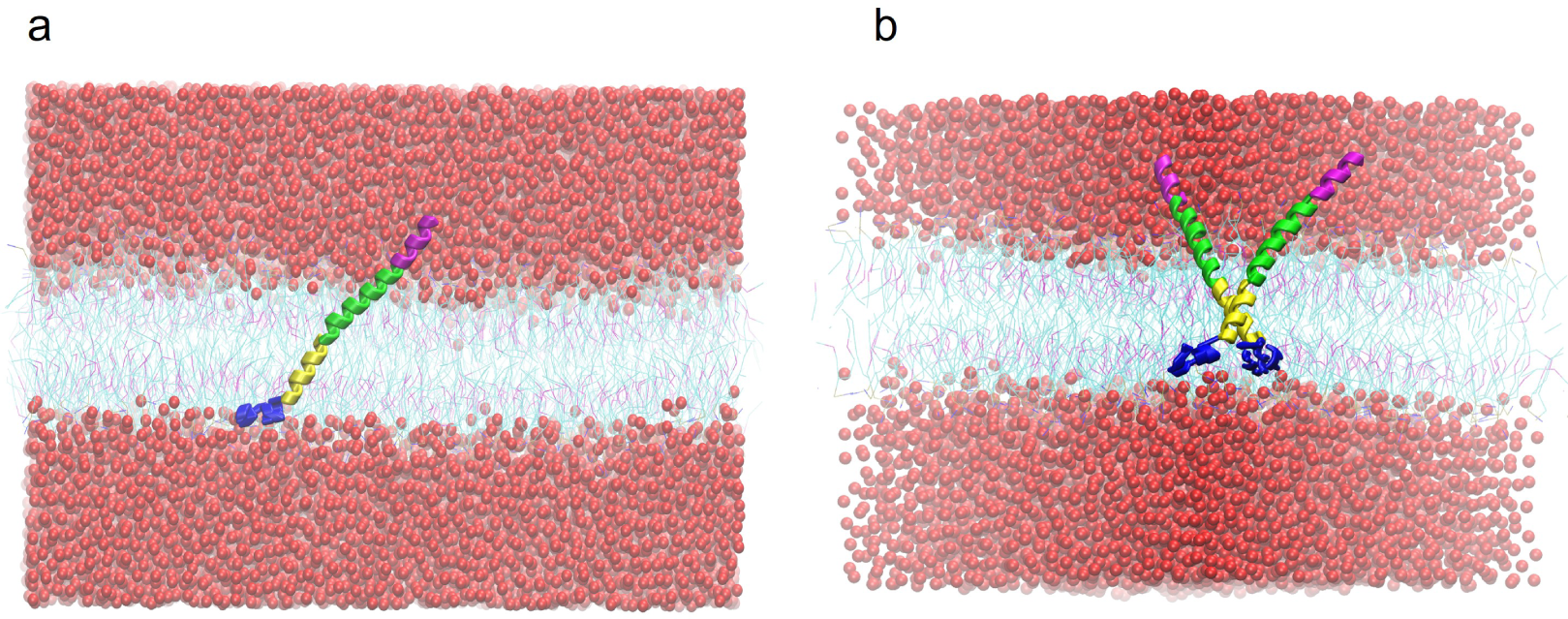
Coarse-grained IRE1^516-571^ in (50%-50%) DOPC-POPC membrane (a) IRE1^516-571^ monomer. The amphipathic portion of the IRE1, IRE1^526-544^(green helix) lies at the interface between the membrane top surface and ER lumen. The rest of IRE1^544-571^(yellow and blue helix) stayed inside the membrane. The purple helix is the IRE1^516-525^ in the ER lumen side. Water represented by red beads. Lipids are represented as violet and cyan lines. (b) IRE1^516-571^ dimer.

### 2.5 Bilayer tension implementation

The surface tension target feature of NAMD was used to regulate bilayer tension in the simulations. The tension, *γ* was applied along the membrane surface. The pressure normal to the membrane surface, P_z_=1 atm. was applied via Langevin piston. The free energy of IRE1^516-571^ dimer dissociation was calculated at four different bilayer tensions, *γ* [15,5,0,-5] pN/nm in NP_z_*γ*T ensemble. Bilayer tension of positive values indicates tension while negative values indicates compression.

### 2.6 Dissociation constant of IRE1 dimerization

The dissociation constant of IRE1 dimerization was calculated via the following equation

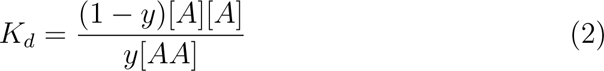

where [*A*] is the concentration of IRE1 monomer, [*AA*] is the concentration of IRE1 dimer, and *y* is the time fraction of IRE1 dimer state. Equation (2) was previously used to calculate *K_d_* for GpA dimers from molecular dynamics simulat [Domański et al., 2017]. The Equation (2) was derived based on the following equilibrium properties: (i) at equilibrium the timed average forward and backward rate of dimerization is equal; and (ii) the time average of the chemical potential of the monomer and dimer at equilibrium are equal. In the dimer state, the concentration of the dimer is, [*AA*] = 1*/σ* and in the monomer state, [*A*] = 2*/σ*, where *σ* represents the area of lipids enclosed by two monomer when they are a furthest apart in the simulation. In this case, *σ*= *πR*^2^*/*4, where R is the distance between the two IRE1 monomer in the farthest bin of the umbrella sampling simulations. By putting these values into Equation (2), we obtain

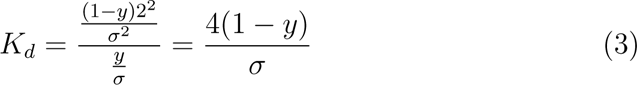

The time fraction of the dimer was determined by the following equation

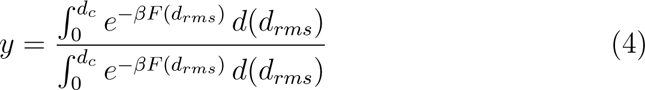

where 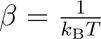, *k*_B_ and T represent Boltzmann’s constant and the absolute temperature, respectively. The critical distance, *d*_c_ separates the region of the dimer state from the monomer state. It was shown previously shown that *y* is relatively insensitive to choice *d*_c_ [Domański et al., 2017]. In this work we use different *d*_c_ values depending on their free energy landscape.

## 3 Results

The free energy landscapes of IRE1^516-571^ dimerization under bilayer tension and compression, calculated by replica-exchange umbrella sampling simulations, are presented in this section. First, to determine how IRE1 monomers are effected by membrane tension we calculate the membrane thickness far from, and close to monomers for each loading condition. Next, we determine how the free energy of dimerization is altered under tension and compression. From the resulting PMF curves, we then estimate the percentage of IRE1^516-571^ dimer at different IRE1 concentrations as a proxy of UPR activation.

### 3.1 Bilayer tension modulates membrane thickness

#### 3.1.1 Bilayer tension decreases the membrane thickness while compression increases the membrane thickness

To determine the effect of membrane tension on bilayer thickness, bilayer tension was applied to a single IRE1^516-571^ monomer embedded in a membrane (50% POPC-50% DOPC). The distance between the phosphate beads of the bottom and top leaflet is defined as the bilayer thickness. Fig. 2 shows membrane thickness variation under the application of tension. With no applied tension, the average membrane thickness is found 3.987 nm (Fig. 2b). In the vicinity of IRE1^516-571^, membrane thickness was depressed to a minimum of 3.720 nm. Hydrophobic mismatch between the lipids and IRE1^516-571^ is the primary cause of this local membrane compression. As expected, applying a compressive force in membrane plane increased the average membrane thickness to 4.044 nm (Fig. 2a). On the other hand, tension applied to the membrane reduces the membrane thickness; tension of 5 pN/nm reduces the average membrane thickness to 3.944 nm and tension 15 pN/nm further reduces the average thickness to 3.853 nm. A plot of bilayer average thickness against bilayer tension is added in Fig. S6. In all cases (both tension and compression) the locally depressed region around IRE1^516-571^ was observed. The observed average membrane thickness variation of the tensed membrane is in accordance with the previous finding of mechanosensitivity of DOPC lipid bilayer [Reddy et al., 2012].

**Fig. 2:**
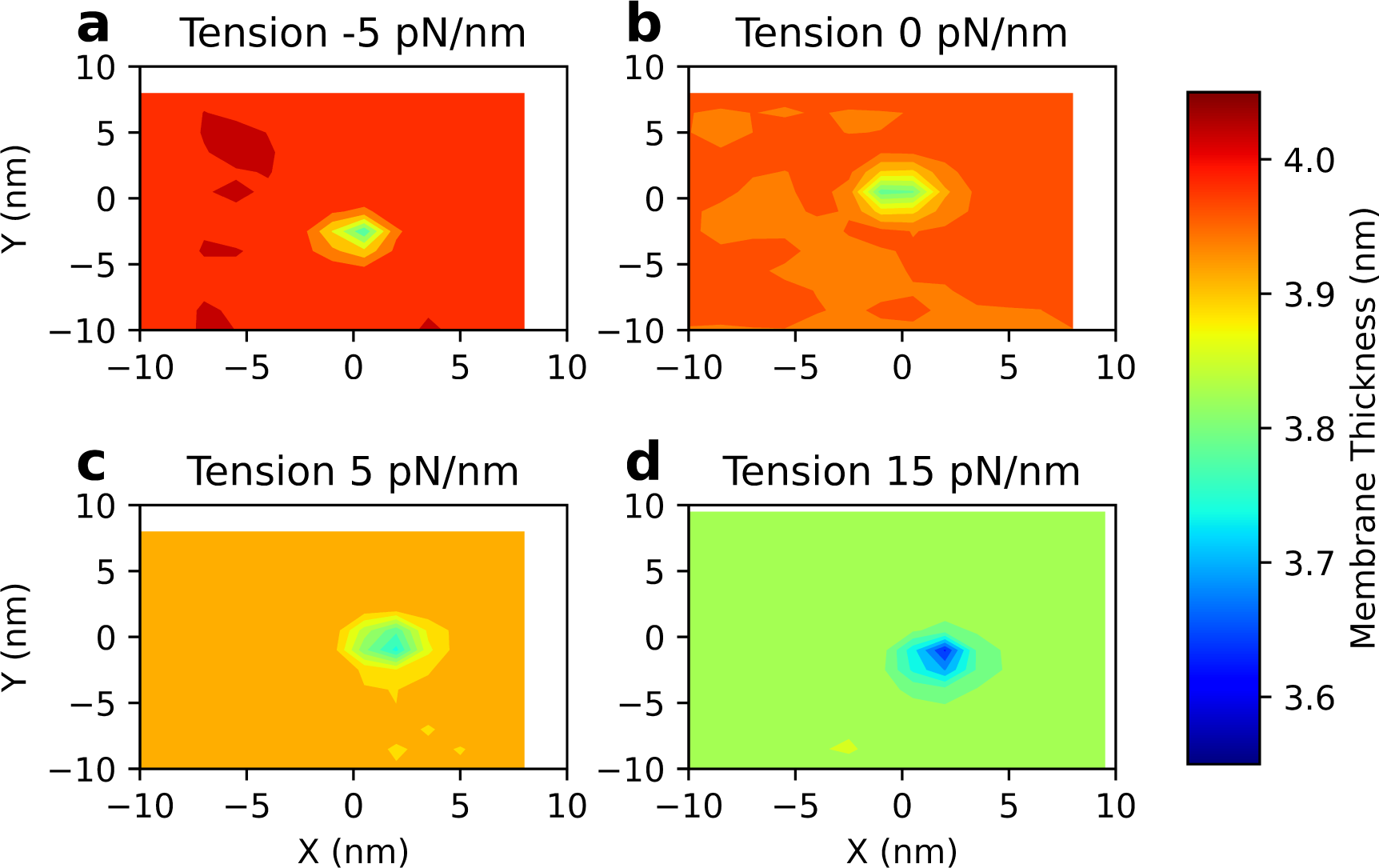
Membrane thickness of a single IRE1^516-571^monomer-(50%DOPC-50%POPC) complex under various bilayer tensions. (a) Compressing the membrane with −5pN/nm increased the average membrane thickness to 4.044 nm. (b) Under zero tension, the average membrane thickness is 3.987 nm. (c) A bilayer tension of 5 pN/nm reduced the average membrane thickness to 3.944 nm. (d) Bilayer tension 15 pN/nm reduced the avergae membrane thickness to 3.853 nm.

#### 3.1.2 Membrane thickness deformation field and bilayer tension

The membrane deformation field, defined as the difference between the local and unperturbed thickness, *u*(*x, y*) = *h*(*x, y*) *− a*, where *h* is the membrane thickness field and *a* is the membrane thickness far from the protein, which is calculated by averaging the thickness over the boundary of the simulation domain. Fig. 3 shows the membrane thickness deformation field around the IRE1^516-571^ monomer for four cases: (a) compression with −5 pN/nm, (b) zero tension, (c) tension 5 pN/nm and, (d) tension 15 pN/nm. In each case, the monomer induces an elliptical depression in the membrane. The degree of depression is more pronounced for each mechanically stressed state.

**Fig. 3:**
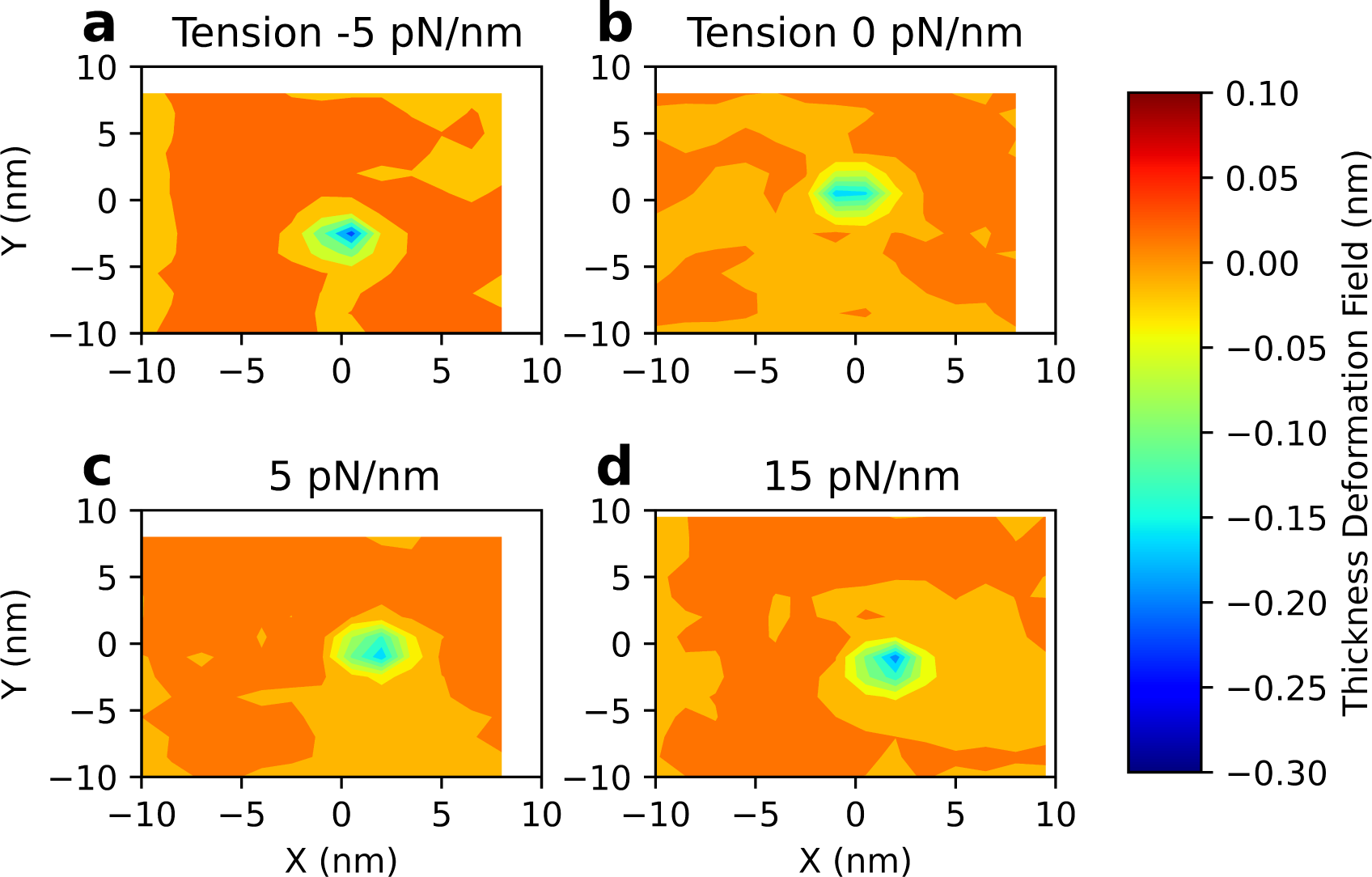
Membrane thickness deformation field in the vicinity of IRE1^516-571^ monomer under application of various bilayer tensions: (a) 5 pN/nm compression, (b) zero tension, (c) 5 pN/nm tension, (d) 15 pN/nm tension.

Next we computed the membrane deformation around IRE1^516-571^ dimers equilibrated at different applied in-plane loads (Fig. 4). The deformations fields remain elliptical and show a similar trend to the monomers, but are more pronounced.

**Fig. 4:**
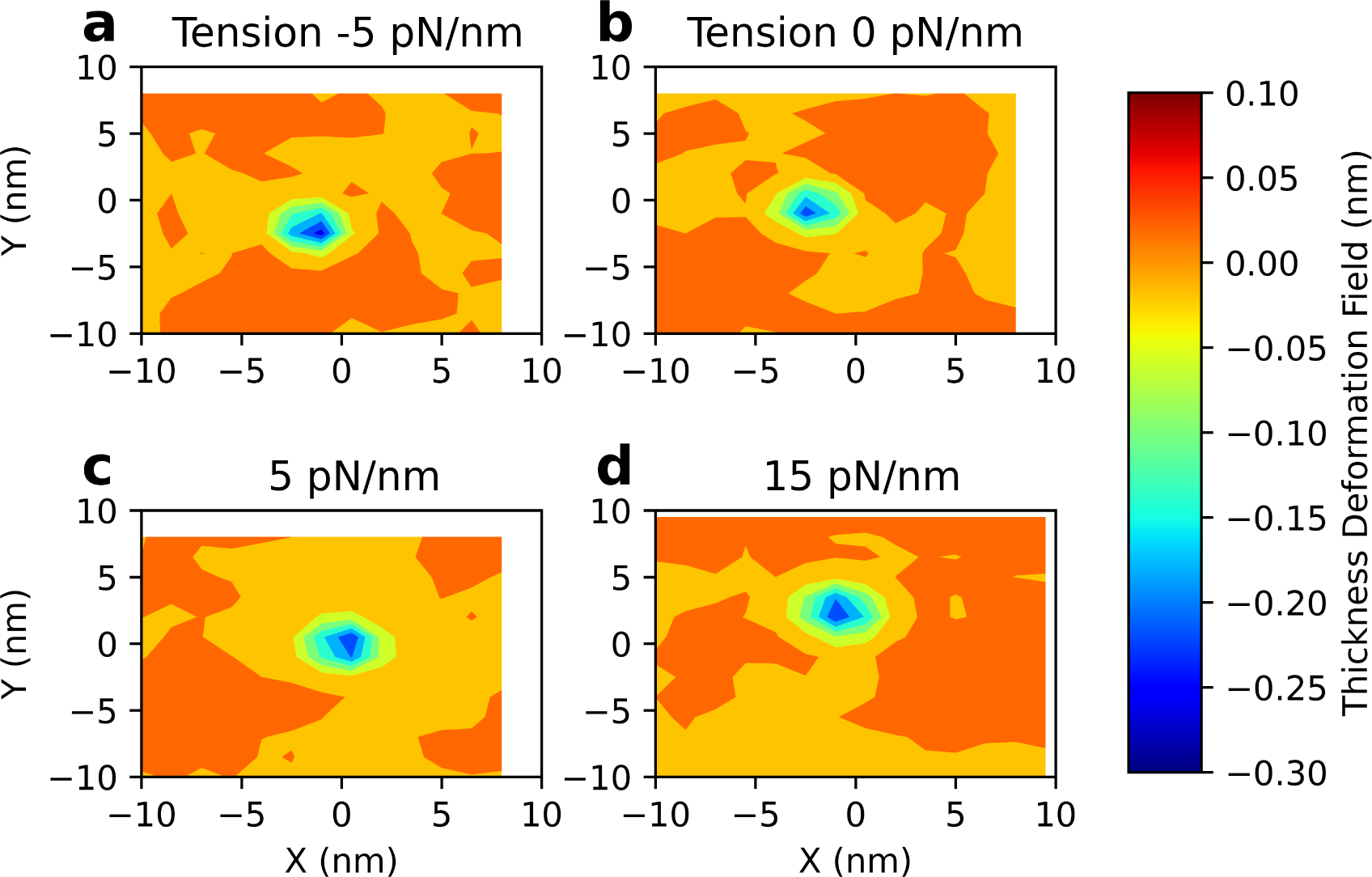
Membrane thickness deformation field in the vicinity of IRE1^516-571^ dimer under application of various bilayer tensions: (a) 5 pN/nm compression, (b) zero tension. (c) 5 pN/nm tension. (d) 15 pN/nm tension.

### 3.2 Bilayer tension and crossing angle

The hydrophobic domain of the IRE1^516-571^ monomer adjusts its orientation in the membrane so that the effective hydrophobic length of IRE1^516-571^ becomes approximately equal to the bilayer thickness. Additionally, IRE1^516-571^ has an amphipathic helix that lies at the interface between ER lumen and the membrane. These properties jointly dictate the orientation of IRE1^516-571^ and the deformation of the membrane around the protein. With no applied tension, IRE1^516-571^ makes an angle of 45*^◦^* with the normal of the membrane surface. Tension of 5 pN/nm reduced membrane thickness to 3.944 nm, which forces IRE1^516-571^ to match its effective hydrophobic length by tilting towards membrane surface. Similarly 15 pN/nm tension increases the tilt angle of the monomer further. On the other hand, membrane compression increases the membrane thickness, reducing the tilt angle. A plot of tilt angle and bilayer tension is shown in Fig. 5. Going from compression to tension, the tilt angle increases. While forming a dimer, two monomer will try to form a X-like structure. The angle between the monomer’s principal axis is defined as the crossing angle. The crossing angle is approximately equal to twice the inclination angle of IRE1 monomer ^516-571^. A schematic of crossing angle is depicted in inset of the Fig. 5. Experimental evidence also showed that two IRE1^516-571^ monomers form an X-like structure resulting in a crossing angle of 90*^◦^* in active dimeric form [Väth et al., 2021]. The simulations were initialized with this active X-like structure of IRE1 dimers. The crossing angle was sustained at 90*^◦^* for no bilayer tension. The average crossing angle under tension was found to be more than 90*^◦^* while in compression, it was below 90*^◦^*. These results show that the membrane-protein dimer complex responds to bilayer tension by both deforming the membrane locally and altering the orientation of the proteins to accommodate the changes in membrane thickness.

**Fig. 5:**
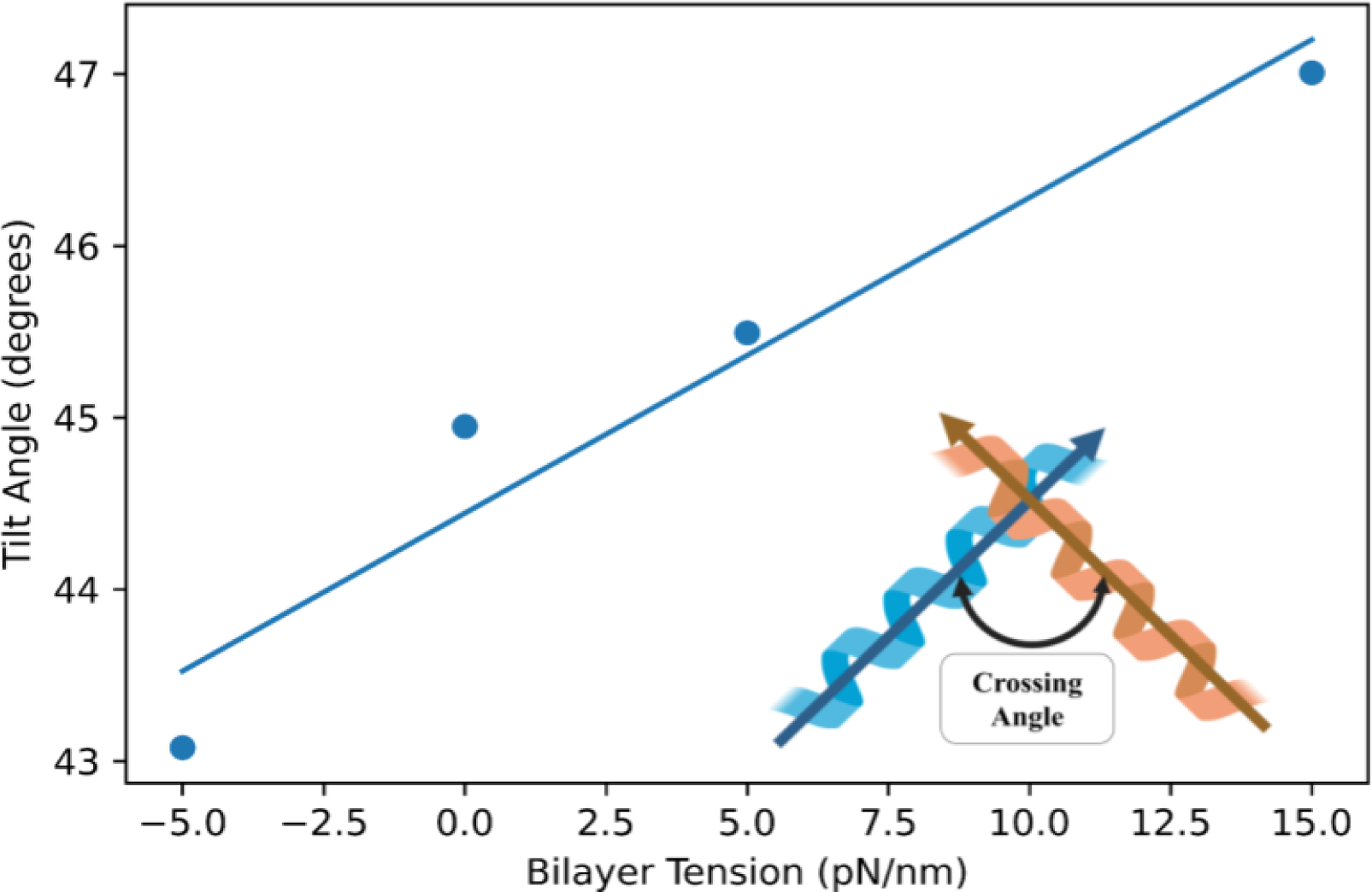
Inclination of IRE1^516-571^ monomer with respect to the normal of membrane surface changes with bilayer tension. Increased bilayer tension increases the inclination towards the membrane surface. In a dimer, IRE1 monomers form a X-like structure and the angle between the two monomers is termed the crossing angle. A schematic of crossing angle is added in the inset.

### 3.3 Variation in bilayer tension favors IRE1 dimerization

Bilayer tension applied to the IRE1^516-571^-membrane complex impacts the free energy of IRE1 dimerization. Fig. 6 shows the potential of mean force for dimer formation under various bilayer tensions. With no externally applied tension, the energy well depth of dimerization is −40.5 kJ/mol. However, when a 5 pN/nm compression is applied, the energy well deepens to −58 kJ/mol, indicating IRE1^516-571^ dimers are more stable in a compressed membrane than in the zero tension state. Similarly, bilayer tension 5 pN/nm deepens the energy minima to −56 kJ/mol. When the tension is further increased to 15 pN/nm, the energy well deepens further to −73 kJ/mol, indicating the most stable IRE1^516-571^ dimer structure among these four cases. These results show that any change in tension, irrespective of direction (tension/compression), favors IRE1^516-571^ dimerization.

**Fig. 6:**
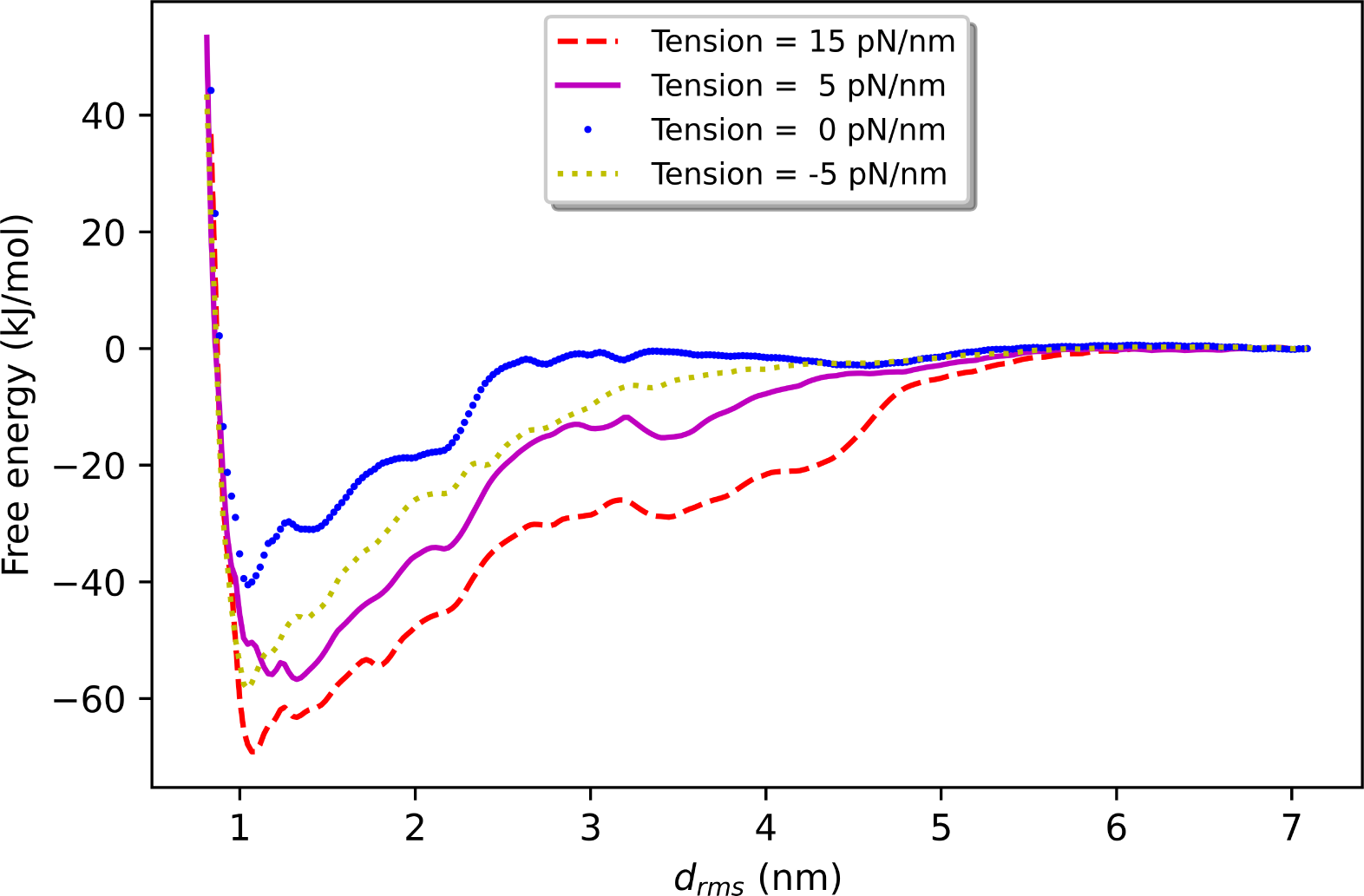
Free energy landscapes of IRE1^516-571^ dimer dissociation as a function *d_rms_* upon application of various bilayer tensions. The blue line corresponds to the free energy of IRE1^516-571^ dimer dissociation at zero bilayer tension, and shows a well depth of −40.5 kJ/mol. The yellow line corresponds to the free energy of IRE1^516-571^ dimer dissociation at compression with a energy well depth of −58 kJ/mol. The magenta line corresponds to the free energy of IRE1^516-571^ dimer dissociation at the tension of 5 pN/nm with a energy well depth of −56 kJ/mol. The red line corresponds to the free energy of IRE1^516-571^ dimer dissociation at tension 15 pN/nm with a energy well depth of −73 kJ/mol. Hence, any perturbation in tension increased the well depth of the free energy curve.

### 3.4 Tension modulates stability of UPR-signalling structure

The calculated PMFs of IRE1^516-571^ dimer dissociation can be used to determine the dissociation constant, *K*_d_ of the IRE1^516-571^ dimer at various bilayer tension. The dissociation constant,*K*_d_ was calculated from equation (2). These *K*_d_ values at different bilayer tension are represented through the binding curves shown in Fig. 7. The binding curves show the ratio of IRE1 in dimer form at different IRE1 concentrations. With no applied tension, the dissociation constant for an IRE1 dimer was found 5.94e8 molecules/nm^2^. *K*_d_ values estimated from simulation differ from *K*_d_ determined by experiment due to simplifications of the coarse-grained model. However, the values reported here are similar to *K*_d_ values of GpA dimerization computed from coarse-grained molecular dynamics simulation [Domański et al., 2017]. 5 pN/nm tension reduced *K*_d_ to 5.49e-11 molecules/nm^2^. As a result, the binding curve under tension shifts left compared to the zero tension state, indicating a higher IRE1 dimer percentage at the same total IRE1 concentration. Similarly, 5 pN/nm compression reduced *K*_d_ to 6.32e-11 molecules/nm^2^, indicating a higher IRE1 dimer concentration compared to zero tension. A bilayer tension of 15 pN/nm further reduced *K*_d_ to 2.32e-13 molecules/nm^2^ exhibiting the highest IRE1 dimer percentage among these four states. Hence, the IRE1 dimer concentration was increases with either tension or compression. The increased IRE1 dimer concentration in tension or compression would favor IRE1 conformations that activate the unfolded protein response.

**Fig. 7:**
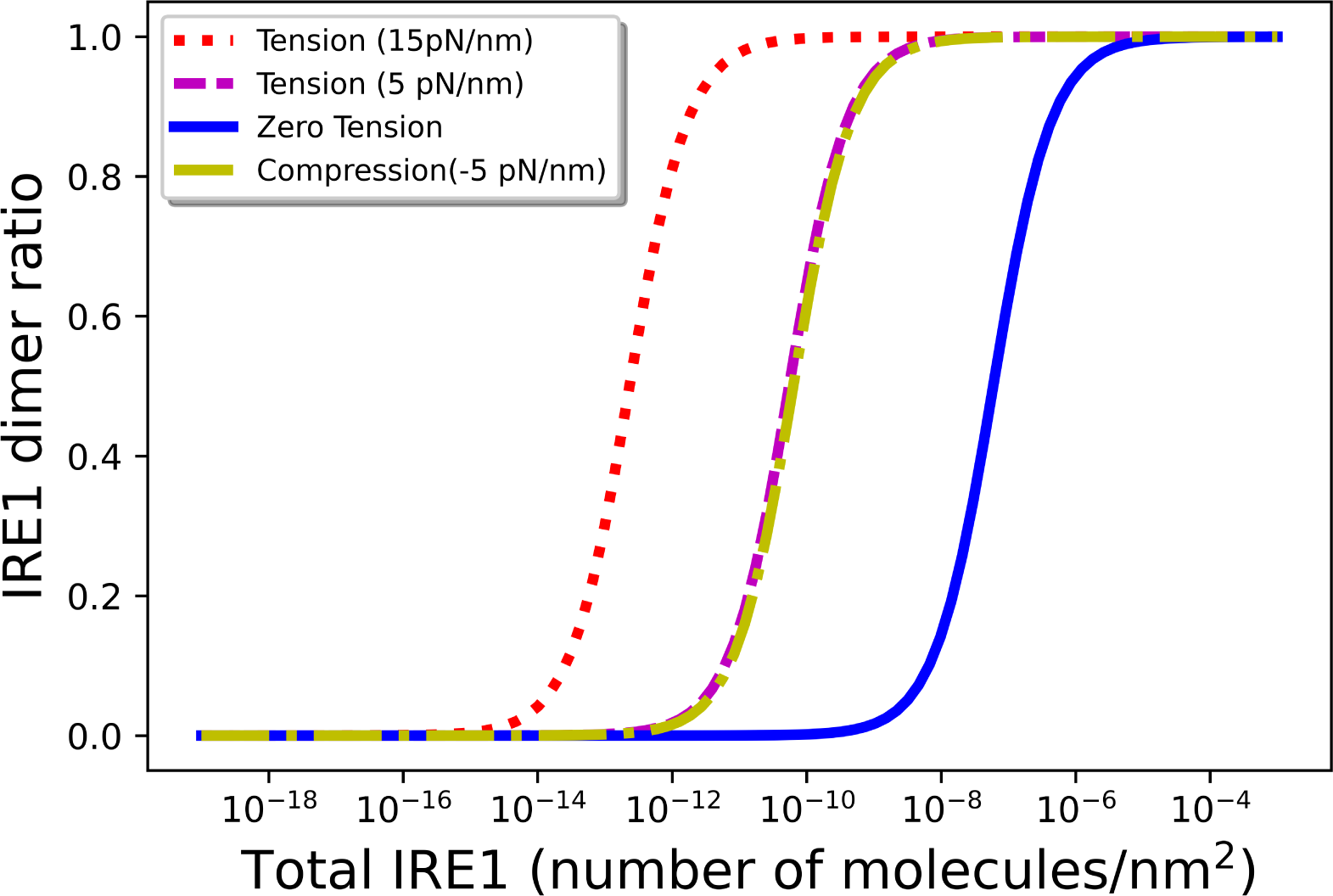
Binding curves of IRE1 dimers under application of various bilayer tensions. Altering bilayer tension changes IRE1 dimer concentration for a particular total concentration of IRE1. When the tension of 5 pN/nm (magenta dashed line) was applied, the binding curve shifted to the left compared to the binding curve of no tension state (blue line). 5 pN/nm (yellow dot-dashed line) compression produced a similar effect. Further application of bilayer tension of 15 pN/nm (red short dashed line) shifts its binding curve further to the left.

## 4 Discussion

As one might expect, in our simulations tension decreased the average membrane thickness, while compression increased the average membrane thickness. Since, in the unperturbed membrane, IRE1 locally reduces the membrane thickness to accommodate hydrophobic mismatch, one might expect externally applied compression to favor dimerization since it would further exacerbate the hydrophobic mismatch of the monomers. Conversely, one might expect tension to favor the monomer state by reducing the hydrophobic mismatch, therefore decreasing the driving force for dimerization. However, our results show that both applied tension and compression enhance the stability of IRE1^516-571^ dimers. How can this be? In the case of tension, the decrease in membrane thickness leads to a decrease in membrane deformation around the IRE1 monomers. However, the energy required for this (smaller) deformation is greater than that of the untensed membrane because the tension prestrains the membrane, effectively increasing the resistance of the system to further deformation. In compression, despite IRE1 attempting to accommodate the thicker membrane by decreasing its tilt angle, the membrane depression is higher than in the untensed membrane due to greater hydrophobic mismatch. This increased deformation also results in increased potential energy of the system.

To see this more clearly, we considered a continuum model of an elastic membrane. The energetic penalty associated with changes in thickness upon insertion of a monomer in a tensed membrane is [Ursell et al., 2007]

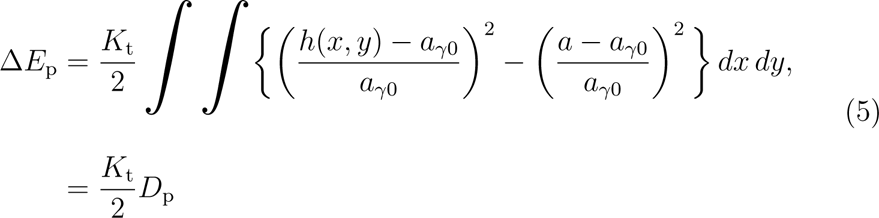

where *a_γ_*_0_ is the unperturbed thickness with no applied tension, and *K*_t_ is the membrane thickness modulus. *D*_p_ is the integrated change in membrane thickness for a loaded membrane upon protein insertion. A plot of Δ*E*_p_ against applied bilayer tension is shown in Fig. 8 (a). Higher Δ*E*_p_ at higher tension indicates more energy is used to deform the membrane around the IRE1^516-571^ monomer when the membrane is under tension. In contrast, under compression, Δ*E*_p_ is lower than that of the zero-tension case. Since lipid bilayers are approximately incompressible [Tosh and Collings, 1986, Seemann and Winter, 2003], membrane thinning is compensated by an increase in the leaflet area. In a compressed membrane, the energy required to increased the lipid area against the applied compression, *γ*, can be estimated as follows [Watson et al., 2013, Bitbol et al., 2012].

**Fig. 8:**
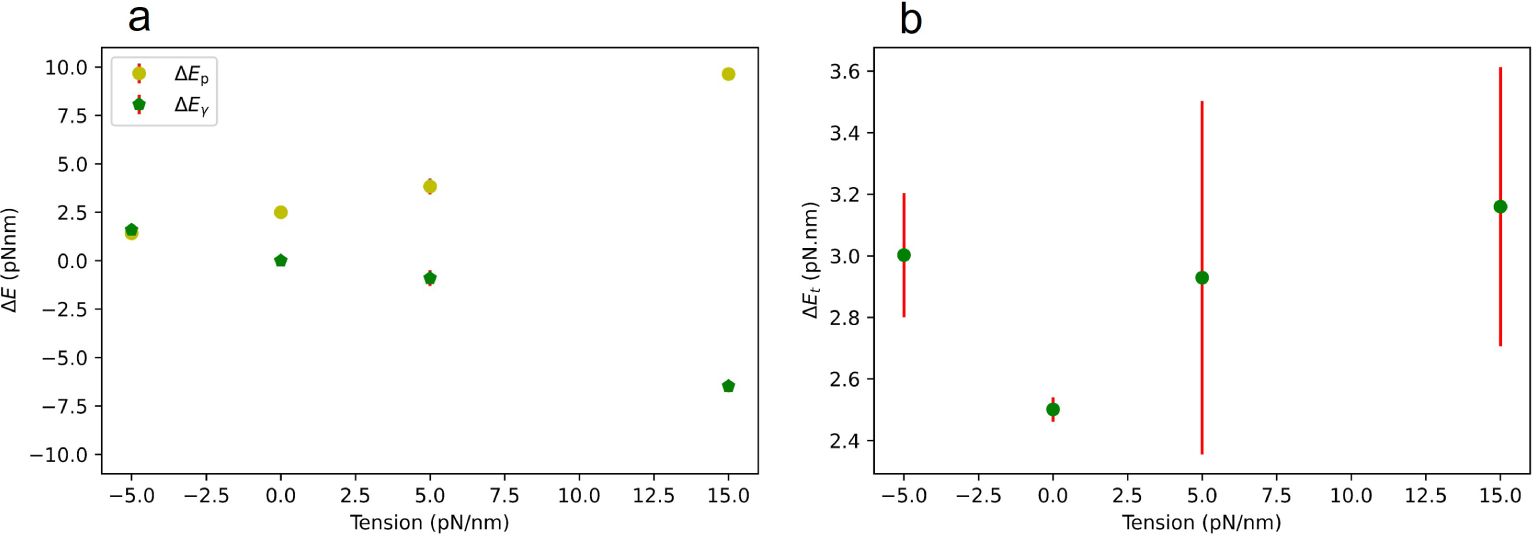
Strain energy contributions due to thickness change and lipid area change upon inserting an IRE1^516-571^ monomer at various bilayer tensions. (a) Δ*E*_p_ and Δ*E_γ_* represents the strain energy contribution due to membrane thickness deformation and lipid area, respectively. Δ*E*_p_ increases with increased applied tension, while Δ*E_γ_* decreases with increased applied tension. (b) Total strain energy change associated with IRE1^516-571^ monomer insertion in ER membrane (50%DOPC-50%POPC) under various bilayer tensions. Δ*E*_t_ at tension zero requires the lowest amount of energy to insert IRE1^516-571^ in the ER. Both tension and compression requires higher energy to insert IRE1^516-571^ in the ER. The calculation of uncertainties is described in Section 3 of Supplementary Material.

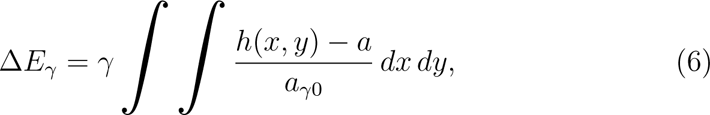

where the change in lipid area is expressed in terms of membrane thickness using the assumption of an incompressible membrane. A plot of Δ*E_γ_* against applied bilayer tension (Fig. 8)(a) shows that, in compression, IRE1^516-571^ requires more energy to change the lipid area than when no load is applied or under tension. The total strain energy due to IRE1^516-571^ insertion in a mechanically stressed ER membrane is the combined effect of both variation of membrane thickness and lipid area, and can be calculated as

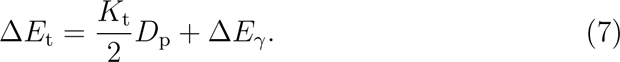

The thickness modulus was found to be 480 pN/nm (see Equation (S3)). Fig. 8 (b) shows Δ*E*_t_ as a function of applied tension, calculated from our simulations. Δ*E*_t_ increases for both tensed and compressed membranes compared to the zero-tension state. Large uncertainties associated with the estimation of Δ*E*_t_ are due to uncertainties in calculating the unperturbed membrane thicknesses in tensed or compressed membranes, similar to other simulations of membrane-protein systems [Argudo et al., 2017]. Δ*E*_t_ at different tensions can be used to compare the energy requirement for IRE1^516-571^ monomer insertion. We see that the continuum model predicts both tension and compression will increase the monomer energy, suggesting a greater driving force for dimerization. Additionally, we see that the driving force for tension is primarily the energy associated with thinning the already-stretched membrane, while the driving force for compression is due to the greater energy needed to create space for the protein in the bilayer. When these contributions are combined, the result is less monomer stability for both compressive and tensile loads.

In cells, IRE1 forms large oligomeric signalling clusters. While our simulations only consider the relative stability of monomers and dimers, we expect that larger signalling clusters would be even more favorable upon mechanical perturbations of the membrane. Since the driving force of dimerization in our study results from decreasing the total deformation energy of the membrane by overlapping the deformation fields of each monomer, it is reasonable to suppose that further decreasing the total area by forming larger oligomeric clusters would be even more favorable. Hence, our study predicts that mechanical perturbations to the ER membrane could lead to the formation of active IRE1 signalling clusters and activation of the UPR in the absence of misfolded protein stress. This is similar to how lipid composition has been shown to regulate the UPR independent of proteotoxic stress.

Despite the extensive simulations used to establish the mechanosensitivity of IRE1^516-571^ in this work, we are unaware of any experimental evidence showing direct activation of the UPR by mechanical forces. Due to computational expense, our study was limited to the transmembrane section of IRE1. Since both the lumenal and cytosolic portions of IRE1 interact with other IRE1 molecules in oligomers, it is possible that full-length IRE1 molecules may behave differently. However, activation of the IRE1 signalling pathway upon lipid compositional changes lends credence to the ability of the trans-membrane section of IRE1 to promote clustering in cell. To confirm the predicted mechanical regulation of the UPR, experimental measurements of full-length IRE1 in tensed membranes are needed.

Sensitivity of IRE1 to membrane tension provides another example of unfolded-protein-independent UPR activation. The tension in the ER membrane would be expected to change as the pressure within the ER changes, for example due to osmotic stress or from an influx of nascent proteins, as well as from cytoskeletal forces applied at anchoring points within the ER membrane. The mechanosensitiviy of IRE1 would allow cells to monitor and respond to these perturbations. Lastly, since the transmembrane domain of PERK is very similar to that of IRE1, and both activate by clustering in the ER membrane, it seems plausible that the PERK signalling pathway may also be sensitive to ER bilayer tension, although further computational and experimental work would be required to confirm this.

## 5 Conclusion

In this study, we analyzed the impact of bilayer tension on IRE1 dimerization through coarse-grained molecular dynamics simulation. Mechanical perturbation of the ER bilayer changes physical properties like the membrane thickness and area per lipid, which impact the free energy of dimerization for a pair of IRE1 monomers. Interestingly, we find that IRE1 dimerization becomes more energetically favorable under both tension and compression as compared to in an unloaded membrane. This manifests as an increase in the ratio of IRE1 dimers to monomers as either tension or compression is applied. From these results, we predict that tension or compression can facilitate UPR activation and that IRE1 may be a mechanosensitive modulator of the ER stress response.

## Acknowledgements

This work was supported by Natural Science and Engineering Research Council of Canada (grant RGPIN-2021-02747).

## References

V. Allen and R. David. The endoplasmic reticulum stress response in the pancreatic *β*-cell. Diabetes, Obesity & Metabolism, 12 Suppl 2(SUPPL. 2): 48–57, 2010. doi: 10.1111/j.1463-1326.2010.01271.x.

D. Argudo, N. P. Bethel, F. V. Marcoline, C. W. Wolgemuth, and M. Grabe. New continuum approaches for determining protein-induced membrane deformations. Biophysical Journal, 112(10):2159–2172, May 2017. doi: 10.1016/j.bpj.2017.03.040.

K. Ben M’barek, D. Ajjaji, A. Chorlay, S. Vanni, L. Forêt, and A. R. Thiam. ER Membrane Phospholipids and Surface Tension Control Cellular Lipid Droplet Formation. Developmental Cell, 41(6):591–604.e7, Jun 2017. doi: 10.1016/j.devcel.2017.05.012.

A.-F. Bitbol, D. Constantin, and J.-B. Fournier. Bilayer elasticity at the nanoscale: The need for new terms. PLOS ONE, 7(11):1–19, Nov 2012. doi: 10.1371/journal.pone.0048306.

I. Braakman and D. N. Hebert. Protein folding in the endoplasmic reticulum. Cold Spring Harbor Perspectives in Biology, 5(5), 2013. doi: 10.1101/cshperspect.a013201.

B. R. Brooks, C. L. Brooks, A. D. Mackerell, L. Nilsson, R. J. Petrella, B. Roux, Y. Won, G. Archontis, C. Bartels, S. Boresch, A. Caflisch, L. Caves, Q. Cui, A. R. Dinner, M. Feig, S. Fischer, J. Gao, M. Hodoscek, W. Im, K. Kuczera, T. Lazaridis, J. Ma, V. Ovchinnikov, E. Paci, R. W. Pastor, C. B. Post, J. Z. Pu, M. Schaefer, B. Tidor, R. M. Venable, H. L. Woodcock, X. Wu, W. Yang, D. M. York, and M. Karplus. CHARMM: The biomolecular simulation program. Journal of Computational Chemistry, 30(10):1545–1614, Jul 2009. doi: 10.1002/jcc.21287.

M. F. Brown. Modulation of rhodopsin function by properties of the membrane bilayer. Chemistry and Physics of Lipids, 73(1-2):159–180, Sep 1994. doi: 10.1016/0009-3084(94)90180-5.

M. Calfon, H. Zeng, F. Urano, J. H. Till, S. R. Hubbard, H. P. Harding, S. G. Clark, and D. Ron. IRE1 couples endoplasmic reticulum load to secretory capacity by processing the XBP-1 mRNA. Nature, 415(6867):92–96, Jan 2002. doi: https://doi.org/10.1038/415092a.

D. E. Clapham. Calcium Signaling. Cell, 131(6):1047–1058, Dec 2007. doi: 10.1016/j.cell.2007.11.028.

J. S. Cox and W. Peter. A novel mechanism for regulating activity of a transcription factor that controls the unfolded protein response. Cell, 87 (3):391–404, Nov 1996. doi: 10.1016/s0092-8674(00)81360-4.

E. Decio L and C. Miriam. ER stress in pancreatic beta cells: the thin red line between adaptation and failure. Science Signaling, 3(110), Feb 2010. doi: 10.1126/scisignal.3110pe7.

J. Domański, G. Hedger, R. B. Best, P. J. Stansfeld, and M. S. Sansom. Convergence and Sampling in Determining Free Energy Landscapes for Membrane Protein Association. The Journal of Physical Chemistry B, 121 (15):3364–3375, 2017. doi: 10.1021/acs.jpcb.6b08445.

J. R. Elliott, D. Needham, J. P. Dilger, and D. A. Haydon. The effects of bilayer thickness and tension on gramicidin single-channel lifetime. Biochimica et Biophysica Acta, 735(1):95–103, Oct 1983. doi: 10.1016/0005-2736(83)90264-x.

P. Fagone and S. Jackowski. Membrane phospholipid synthesis and endoplasmic reticulum function. Journal of Lipid Research, 50(SUPPL.):S311, Apr 2009. doi: 10.1194/jlr.R800049-jlr200.

G. Fiorin, M. L. Klein, and J. Hénin. Using collective variables to drive molecular dynamics simulations. Molecular Physics, 111(22-23):3345–3362, 2013. doi: 10.1080/00268976.2013.813594.

B. M. Gardner, D. Pincus, K. Gotthardt, C. M. Gallagher, and P. Walter. Endoplasmic reticulum stress sensing in the unfolded protein response. Cold Spring Harbor Perspectives in Biology, 5(3):a013169–a013169, Feb 2013. doi: 10.1101/cshperspect.a013169.

A. Grossfield. Wham: the weighted histogram analysis method, version 2.0.11.

R. Guixà-González, I. Rodriguez-Espigares, J. M. Ramírez-Anguita, P. Carrió-Gaspar, H. Martinez-Seara, T. Giorgino, and J. Selent. MEM-BPLUGIN: studying membrane complexity in VMD. Bioinformatics, 30 (10):1478–1480, Jan 2014. doi: 10.1093/bioinformatics/btu037.

H. Gökhan S. Endoplasmic reticulum stress and the inflammatory basis of metabolic disease. Cell, 140(6):900–917, 2010. doi: 10.1016/j.cell.2010.02.034.

K. Halbleib, K. Pesek, R. Covino, H. F. Hofbauer, D. Wunnicke, I. Hänelt, G. Hummer, and R. Ernst. Activation of the Unfolded Protein Response by Lipid Bilayer Stress. Molecular Cell, 67(4):673–684.e8, 2017. doi: 10.1016/j.molcel.2017.06.012.

D. N. Hebert, S. C. Garman, and M. Molinari. The glycan code of the endoplasmic reticulum: Asparagine-linked carbohydrates as protein maturation and quality-control tags. Trends in Cell Biology, 15(7):364–370, Jul 2005. doi: 10.1016/j.tcb.2005.05.007.

C. Hetz. The unfolded protein response: controlling cell fate decisions under ER stress and beyond. Nature Reviews Molecular Cell Biology 2012 13:2, 13(2):89–102, Jan 2012. doi: 10.1038/nrm3270.

C. Hetz and S. Saxena. ER stress and the unfolded protein response in neurodegeneration. Nature Reviews Neurology, 13(8):477–491, Aug 2017. doi: 10.1038/nrneurol.2017.99.

P. C. Hsu, B. M. Bruininks, D. Jefferies, P. Cesar Telles de Souza, J. Lee, D. S. Patel, S. J. Marrink, Y. Qi, S. Khalid, and W. Im. Charmmgui martini maker for modeling and simulation of complex bacterial membranes with lipopolysaccharides. Journal of Computational Chemistry,, 38(27):2354– 2363, 2017. doi: 10.1002/jcc.24895.

W. Humphrey, A. Dalke, and K. Schulten. VMD – Visual Molecular Dynamics. Journal of Molecular Graphics, 14:33–38, 1996.

S. Jo, T. Kim, V. G. Iyer, and W. Im. CHARMM-GUI: A web-based graphical user interface for CHARMM. Journal of Computational Chemistry, 29 (11):1859–1865, Aug 2008. doi: 10.1002/jcc.20945.

S. Jo, J. B. Lim, J. B. Klauda, and W. Im. CHARMM-GUI membrane builder for mixed bilayers and its application to yeast membranes. Biophysical Journal, 97(1):50–58, Jul 2009. doi: 10.1016/j.bpj.2009.04.013.

J. A. Killian and T. K. Nyholm. Peptides in lipid bilayers: the power of simple models. Current Opinion in Structural Biology, 16(4):473–479, Aug 2006. doi: 10.1016/j.sbi.2006.06.007.

J. B. Klauda, R. M. Venable, J. A. Freites, J. W. O’Connor, D. J. Tobias, C. Mondragon-Ramirez, I. Vorobyov, A. D. MacKerell, and R. W. Pastor. Update of the CHARMM all-atom additive force field for lipids: Validation on six lipid types. The Journal of Physical Chemistry. B, 114(23):7830, Jun 2010. doi: 10.1021/jp101759q.

J. Lee, X. Cheng, J. M. Swails, M. S. Yeom, P. K. Eastman, J. A. Lemkul, S. Wei, J. Buckner, J. C. Jeong, Y. Qi, S. Jo, V. S. Pande, D. A. Case, C. L. Brooks, A. D. MacKerell, J. B. Klauda, and W. Im. CHARMM-GUI Input Generator for NAMD, GROMACS, AMBER, OpenMM, and CHARMM/OpenMM Simulations Using the CHARMM36 Additive Force Field. Journal of Chemical Theory and Computation, 12(1):405–413, Jan 2016. doi: 10.1021/acs.jctc.5b00935.

J. Lee, D. S. Patel, J. Ståhle, S. J. Park, N. R. Kern, S. Kim, J. Lee, X. Cheng, M. A. Valvano, O. Holst, Y. A. Knirel, Y. Qi, S. Jo, J. B. Klauda, G. Widmalm, and W. Im. CHARMM-GUI Membrane Builder for Complex Biological Membrane Simulations with Glycolipids and Lipoglycans. Journal of Chemical Theory and Computation, 15(1):775–786, Jan 2019. doi: 10.1021/acs.jctc.8b01066.

S. J. Marrink, H. J. Risselada, S. Yefimov, D. P. Tieleman, and A. H. De Vries. The MARTINI force field: Coarse grained model for biomolecular simulations. The Journal of Physical Chemistry B, 111(27):7812–7824, 2007. doi: 10.1021/jp071097f.

O. G. Mouritsen and M. Bloom. Mattress model of lipid-protein interactions in membranes. Biophysical Journal, 46(2):141–153, Aug 1984. doi: 10.1016/S0006-3495(84)84007-2.

H. S. Muddana, R. R. Gullapalli, E. Manias, and P. J. Butler. Atomistic simulation of lipid and DiI dynamics in membrane bilayers under tension. Physical Chemistry Chemical Physics, 13(4):1368, Jan 2011. doi: 10.1039/c0cp00430h.

E. Perozo, A. Kloda, D. M. Cortes, and B. Martinac. Physical principles underlying the transduction of bilayer deformation forces during mechanosensitive channel gating. Nature Structural Biology, 9(9):696–703, 2002. doi: 10.1038/nsb827.

J. C. Phillips, D. J. Hardy, J. D. Maia, J. E. Stone, J. V. Ribeiro, R. C. Bernardi, R. Buch, G. Fiorin, J. Hénin, W. Jiang, R. McGreevy, M. C. Melo, B. K. Radak, R. D. Skeel, A. Singharoy, Y. Wang, B. Roux, A. Aksimentiev, Z. Luthey-Schulten, L. V. Kalé, K. Schulten, C. Chipot, and E. Tajkhorshid. Scalable molecular dynamics on CPU and GPU architectures with NAMD. Journal of Chemical Physics, 153(4):044130, Jul 2020. doi: 10.1063/5.0014475.

B. Pontes, P. Monzo, and N. C. Gauthier. Membrane tension: A challenging but universal physical parameter in cell biology. Seminars in Cell & Developmental Biology, 71:30–41, 2017. doi: https://doi.org/10.1016/j.semcdb.2017.08.030.

T. A. Rapoport. Protein translocation across the eukaryotic endoplasmic reticulum and bacterial plasma membranes. Nature, 450(7170):663–669, Nov 2007. doi: 10.1038/nature06384.

W. Rawicz, K. C. Olbrich, T. McIntosh, D. Needham, and E. A. Evans. Effect of chain length and unsaturation on elasticity of lipid bilayers. Biophysical Journal, 79(1):328, 2000. doi: 10.1016/s0006-3495(00)76295-3.

A. S. Reddy, D. T. Warshaviak, and M. Chachisvilis. Effect of membrane tension on the physical properties of dopc lipid bilayer membrane. Biochimica et Biophysica Acta (BBA) - Biomembranes, 1818(9):2271–2281, 2012. doi: https://doi.org/10.1016/j.bbamem.2012.05.006.

D. W. Reid and C. V. Nicchitta. Diversity and selectivity in mRNA translation on the endoplasmic reticulum. Nature Reviews Molecular Cell Biology, 16(4):221–231, Apr 2015. doi: 10.1038/nrm3958.

D. Ron and P. Walter. Signal integration in the endoplasmic reticulum unfolded protein response. Nature Reviews Molecular Cell Biology, 8(7):519– 529, Jul 2007. doi: 10.1038/nrm2199.

U. Schmidt, G. Guigas, and M. Weiss. Cluster formation of transmembrane proteins due to hydrophobic mismatching. Physical Review Letters, 101 (12):1–4, 2008. doi: 10.1103/physrevlett.101.128104.

H. Seemann and R. Winter. Volumetric properties, compressibilities and volume fluctuations in phospholipid-cholesterol bilayers. Zeitschrift für Physikalische Chemie, 217(7):831–846, 2003. doi: doi:10.1524/zpch.217.7.831.20388.

M. M. Sperotto and O. G. Mouritsen. Dependence of lipid membrane phase transition temperature on the mismatch of protein and lipid hydrophobic thickness. European Biophysics Journal, 16(1):1–10, May 1988. doi: 10.1007/bf00255320.

M. M. Sperotto, J. H. Ipsen, and O. G. Mouritsen. Theory of protein-induced lateral phase separation in lipid membranes. Cell Biophysics, 14(1):79–95, Feb 1989. doi: 10.1007/bf02797393.

E. Szegezdi, S. E. Logue, A. M. Gorman, and A. Samali. Mediators of endoplasmic reticulum stress-induced apoptosis. EMBO Reports, 7(9):880, Sep 2006. doi: 10.1038/sj.embor.7400779.

R. E. Tosh and P. J. Collings. High pressure volumetric measurements in dipalmitoylphosphatidylcholine bilayers. Biochimica et Biophysica Acta (BBA) - Biomembranes, 859(1):10–14, 1986. ISSN 0005-2736. doi: https://doi.org/10.1016/0005-2736(86)90312-3.

T. Ursell, K. C. Huang, E. Peterson, and R. Phillips. Cooperative gating and spatial organization of membrane proteins through elastic interactions. PLoS Computational Biology, 3(5):e81, May 2007. doi: 10.1371/journal.pcbi.0030081.

K. Väth, C. Mattes, J. Reinhard, R. Covino, H. Stumpf, G. Hummer, and R. Ernst. Cysteine cross-linking in native membranes establishes the trans-membrane architecture of Ire1. Journal of Cell Biology, 220(8), 2021. doi: 10.1083/jcb.202011078.

M. C. Watson, A. Morriss-Andrews, P. M. Welch, and F. L. H. Brown. Thermal fluctuations in shape, thickness, and molecular orientation in lipid bilayers. II. Finite surface tensions. The Journal of Chemical Physics, 139 (8), Aug 2013. ISSN 0021-9606. doi: 10.1063/1.4818530.084706.

L. M. Westrate, J. E. Lee, W. A. Prinz, and G. K. Voeltz. Form follows function: The importance of endoplasmic reticulum shape. Annual Review of Biochemistry, 84:791–811, Jun 2015. doi: 10.1146/annurev-biochem-072711-163501.

L. Xuemei, Z. Kezhong, and L. Zihai. Unfolded protein response in cancer: the physician’s perspective. Journal of Hematology & Oncology, 4, 2011. doi: 10.1186/1756-8722-4-8.

H. Yoshida, T. Matsui, A. Yamamoto, T. Okada, and K. Mori. XBP1 mRNA is induced by ATF6 and spliced by IRE1 in response to ER stress to produce a highly active transcription factor. Cell, 107(7):881–891, Dec 2001. doi: 10.1016/S0092-8674(01)00611-0.

