## Supplementary Material for "Bilayer tension-induced clustering of the UPR sensor IRE1"

**Md Z. Hossain & W. Stroberg**

This supplementary material consists of three sections. Section 1 demonstrates the convergence test of replica exchange umbrella simulations. This provides evidence that simulations were converged. The last Section 2 shows how the geometrical shape of ER changed upon application of bilayer tension. In the last Section 3, we discussed how some numerical parameters like thickness modulus, $K_{\mathrm{t}}$, $\Delta E_{\mathrm{p}}$ and $\Delta E_{\gamma}$ were estimated from trajectories of IRE1$^{516\text{-}571}$ monomer in 50% DOPC- 50%POPC composition.

#### 1 Convergence analysis

For each tension case, a convergence test was done in the following method; the longest simulation is taken as the ground truth or reference, $T_{\mathrm{total}}$ and the total simulation was divided into several segments, $T_{\mathrm{i}}$, where i represents a segment of total simulation trajectory. The root mean square error of free energy profile of IRE1 dimer dissociation computed by $i^{\mathrm{th}}$ simulation compared to reference free energy profile of IRE1 dimer dissociation($T_{\mathrm{total}}$) is calculated by following equation:

$$RMSE_i = \sqrt{\sum_{j=5}^{71} \frac{F_t(j) - F_i(j)}{n}} \tag{S1}$$

Where n represents the number of reaction coordinates in the free energy landscape. $F_{\mathrm{t}}(j)$ and $F_{\mathrm{i}}(j)$ represents the free energy of the total simulation and free energy of the segment i at reaction coordinate j in terms of kJ/mol-K respectively. Plot of $RMSE_{\mathrm{i}}$ against the independent segments of simulation shows that increasing the simulation time reduces the $RMSE_{\mathrm{i}}$ because of increased samplings. Fig. S1 shows a plot of RMSE against different simulation timescales for bilayer tension 15 pN/nm. Panel S1(b) shows free energy landscapes of three independent 1

$\mu$s simulations compared to the free energy profile of a total 3 $\mu$s simulation. Similarly, Fig S2, S3, and S4 show the convergence of bilayer tension 5, 0, and -5 pN/nm, respectively.

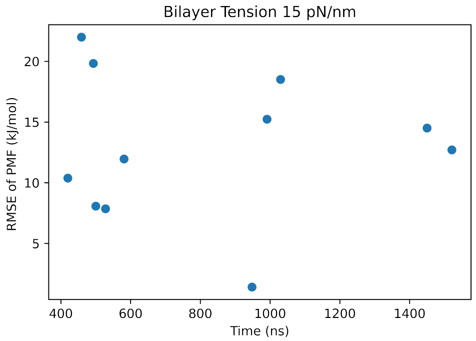


(a) RMSE values of free energy profiles of independent simulations with respect to total simulation.

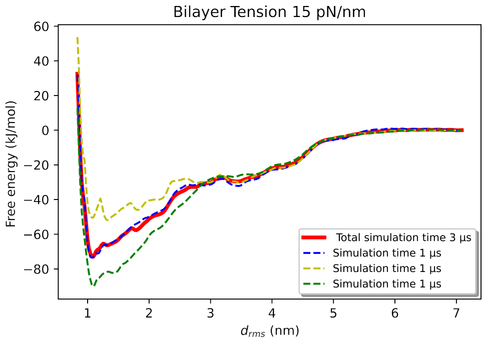


(b) Free energy profiles of three independent 1 $\mu$s simulations compared to reference 3 $\mu$s simulation.

Supplementary Figure S1: Convergence analysis of free energy landscape of IRE1 dimer dissociation at bilayer tension 15 pN/nm.

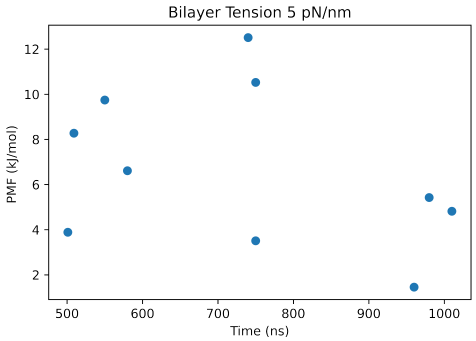


(a) RMSE values of free energy profiles of independent simulations with respect to total simulation.

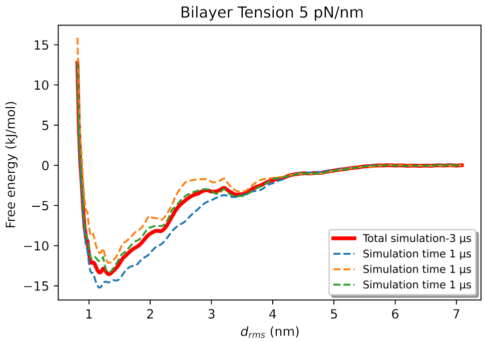


(b) Free energy profiles of three independent 1 $\mu$s simulations compared to reference 3 $\mu$s simulation.

Supplementary Figure S2: Convergence analysis of free energy landscape of IRE1 dimer dissociation at bilayer tension 5 pN/nm.

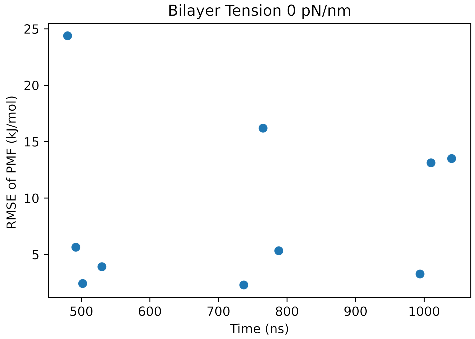


(a) RMSE values of free energy profiles of independent simulations with respect to total simulation.

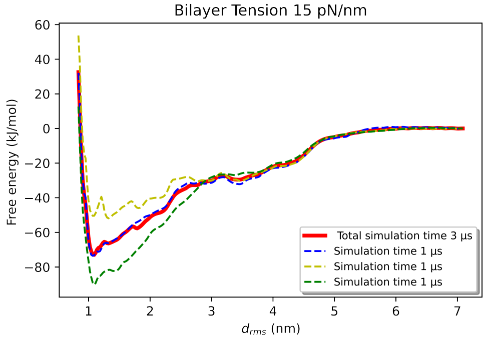


(b) Free energy profiles of three independent 1 $\mu$s simulations compared to reference 3 $\mu$s simulation.

Supplementary Figure S3: Convergence analysis of free energy landscape of IRE1 dimer dissociation at bilayer tension 0 pN/nm.

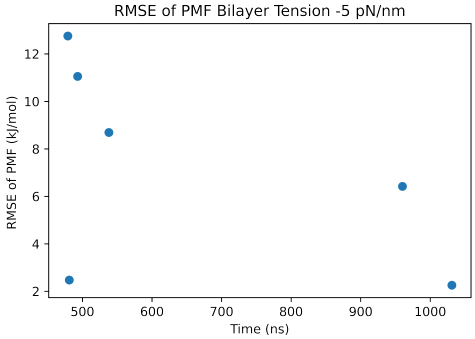


(a) RMSE values of free energy profiles of independent simulations with respect to total simulation at bilayer tension -5 pN/nm.

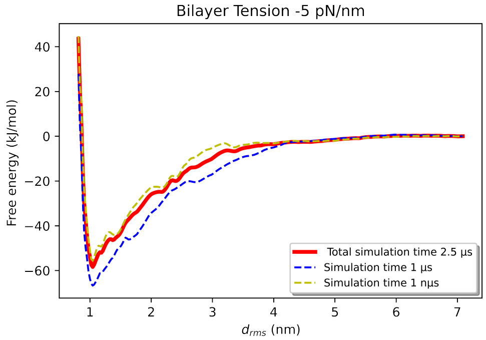


(b) Free energy profiles of two independent 1 $\mu$s simulations compared to reference 2.5 $\mu$s simulation at bilayer tension -5 pN/nm.

Supplementary Figure S4: Convergence analysis of free energy landscape of IRE1 dimer dissociation at bilayer tension -5 pN/nm.

### 2 Geometric parameter of ER

#### 2.1 Area per lipid

Tension applied across the membrane surface stretches the membrane, increasing the surface area. On the other hand, compressing the ER reduced the area. Area per lipid is defined as

$$A_l = \frac{A_{xy}}{N} = \frac{l_x * l_y}{N} \tag{S2}$$

Where $A_\mathrm{l}$ and A represents the area per lipid and membrane surface area, respectively. $l_\mathrm{x}$, $l_\mathrm{y}$ are the x-y dimensions of the membrane surface, A. N is the total number of lipids. Since bilayer tension changes the membrane surface area, it also changes the area per lipid. Since applied tension increases membrane surface area, the area per lipid is also increased. For bilayer tension of 5 pN/nm, $A_\mathrm{l}$ was increased to 66.5 Å$^2$ from 65.5 Å$^2$ of bilayer tension zero. The tension of 15 pN/nm further increased the $A_\mathrm{l}$ to 69 Å$^2$. Compression of -5 pN/nm by decreasing membrane surface area also decreased $A_\mathrm{l}$ to 64.25 Å$^2$. By fitting the values of $A_\mathrm{l}$ with respect to bilayer tension, we find a linear relationship between them (Fig. S5).

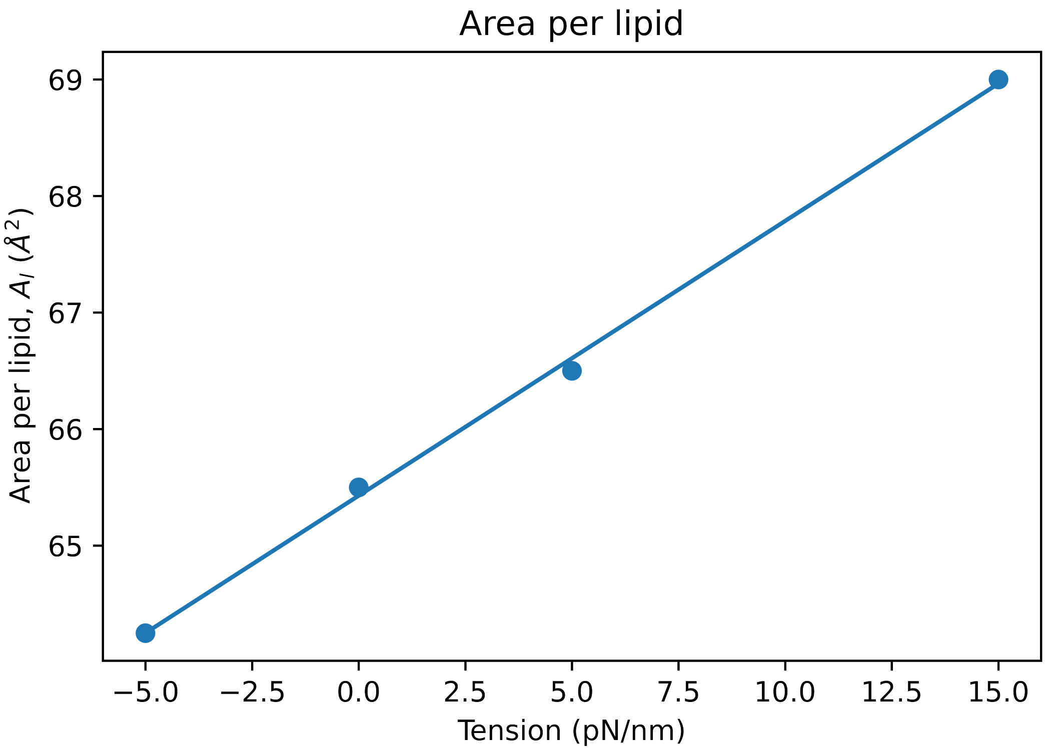


Supplementary Figure S5: Area per lipid shows a linear relationship with bilayer tension. For example, going from compression of -5 pN/nm to tension of 15 pN/nm increased the area per lipid from 64.5 Å$^2$ to 69 Å$^2$.

##### 2.2 Membrane thickness

Membrane thickness is defined as the distance between the lipid heads of two leaflets. For a coarse-grained membrane, it is defined as the distance between PO4 beads of two leaflets. Bilayer tension increases the membrane surface. In doing so, lipid tails are more compressed along their lengths. As a result, membrane thickness reduces under the application of tension. In no tension state, the average membrane thickness is 3.988 nm. The 5 and 15 pN/nm tension reduced the membrane thickness to 3.94 nm and 3.85 nm, respectively. Compression reduced the membrane surface and impacted the lipid tails to extend along its length. So, -5 pN/nm compression increased the membrane thickness to 4.04 nm. Fig. S6 shows a plot of membrane thickness against bilayer tension. One standard deviation of average membrane thickness is shown as the standard error. This plot shows a linear relationship between membrane thickness and bilayer tension. Fig. S5 and Fig. S6 indicate that ER shape was changed under tension/compression.

#### 3 Numerical parameters

##### 3.1 Estimation of thickness modulus

The thickness modulus of the lipid composition, $K_{\mathrm{t}}$ was estimated by following equation[Watson et al., 2013, Bitbol et al., 2012]

$$a_\gamma = a_{\gamma 0}(1 - \gamma/K_t) \tag{S3}$$

where $\gamma$ is the applied bilayer tension and $a_\gamma$ and $a_{\gamma 0}$ is the unperturbed bilayer thickness at the bilayer tension $\gamma$ and 0 respectively. To calculate the $a_\gamma$ and $a_{\gamma 0}$ , we have divided the entire 3 $\mu$s simulation trajectory into independent sections of 1 ns simulation and calculated the far field average of membrane thickness with the help of VMD plugin MEMB and python codes. By plotting the $a_\gamma$ and $a_{\gamma 0}$ against the applied bilayer tension $\gamma$, we estimated the $K_{\mathrm{t}}$ to be 480 pN/nm.

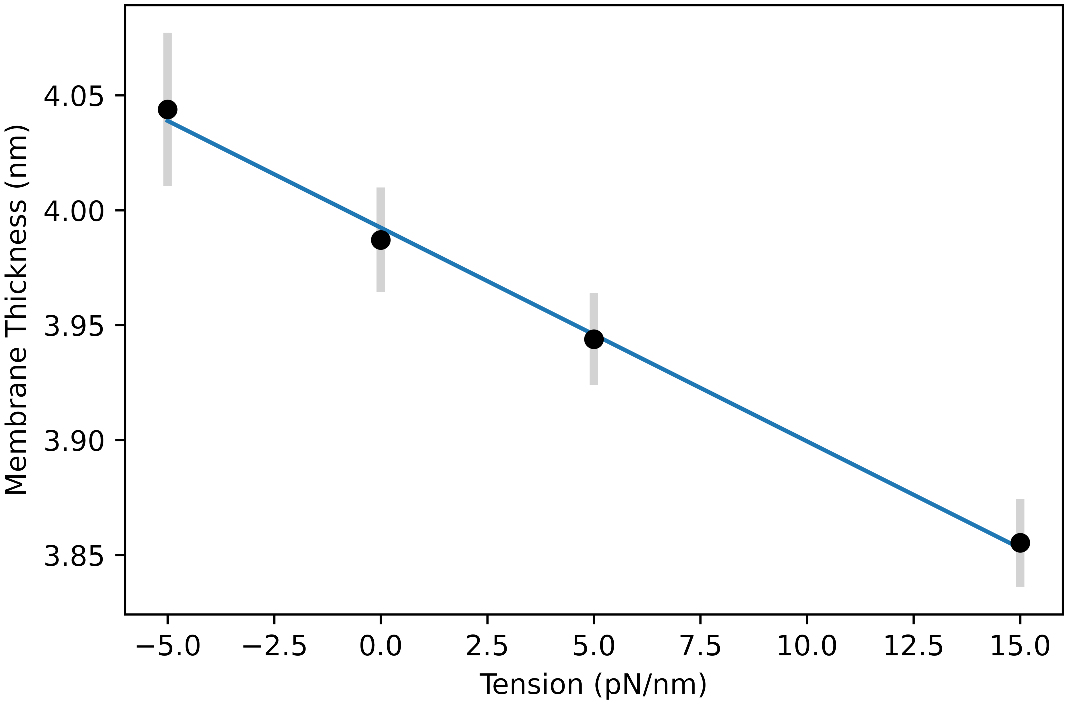


Supplementary Figure S6: Membrane thickness varied linearly with applied bilayer tension. Going from compression of -5 pN/nm to tension of 15 pN/nm decreased the membrane thickness from 4.04 nm to 3.85 nm.

##### 3.2 Calculation of $\Delta E_\mathbf{p}$

For calculation of $\Delta E_\mathrm{p}$, following equation was used

$$\Delta E_\mathrm{p} = \frac{K_t}{2} \int \int \left[ \left\{ \left( \frac{h(x,y) - a_{\gamma 0}}{a_{\gamma 0}} \right)^2 - \left( \frac{a - a_{\gamma 0}}{a_{\gamma 0}} \right)^2 \right\} \right] dx\, dy \qquad \text{(S4)}$$

First, we have determined the unperturbed bilayer thickness at zero tension, $a_{\gamma 0}$. To calculate unperturbed, we have used membrane thickness plot of three independent 1 $\mu$s simulation trajectories of single IRE1$^{516\text{-}571}$ monomer into 50% DOPC-50%POPC lipid. The unperturbed thickness of tensed/compressed ER was also calculated by analyzing three independent 1 $\mu$s simulations totaling 3 $\mu$s simulations. For each 1 $\mu$s simulation trajectory, membrane thickness plots were produced via the MEMB plugin of VMD. $a$ for each tension case was calculated by averaging bilayer thickness far from the position of IRE1$^{516\text{-}571}$ inclusion. The heat map of membrane thickness of 1 $\mu$s trajectory under bilayer tension 5 pN/nm is shown in Fig. S7a. The unperturbed membrane thickness was calculated by averaging the thickness over the surface, excluding the area influenced by the inclusion of IRE1$^{516\text{-}571}$. The membrane area depressed by IRE1$^{516\text{-}571}$ is highlighted by a blue circle in Fig. S7a. Next, membrane deformation, $u(x,y) = h(x,y) - a$ is shown in Fig. S7b. For uncertainty calculation of $\Delta E_{\rm p}$, $\Delta E_{\rm p}$ can be arranged as follows

$$\begin{aligned}\Delta E_{\rm p} =& \frac{K_t}{2}\int\int\left[\left\{\left(\frac{h(x,y)-a_{\gamma 0}}{a_{\gamma 0}}\right)^2 - \left(\frac{a-a_{\gamma 0}}{a_{\gamma 0}}\right)^2\right\}\right] dx\, dy \\ =& \frac{K_t}{2}\int\int\left(\frac{h(x,y)-a_{\gamma 0}}{a_{\gamma 0}}\right)^2 dx\, dy - \frac{K_t}{2}\int\int\left(\frac{a-a_{\gamma 0}}{a_{\gamma 0}}\right)^2 dx\, dy \\ =& \Delta E_{\rm p1} - \Delta E_{\rm p2}\end{aligned} \tag{S5}$$

Then, the uncertainty of $\Delta E_{\rm p}$ was estimated by following equation

$$\Delta\Delta D_p = \sqrt{\left(\frac{\partial \Delta E_{p1}}{\partial a_{\gamma 0}}\Delta a_{\gamma 0}\right)^2 + \left(\frac{\partial \Delta E_{p2}}{\partial a_{\gamma 0}}\Delta a_{\gamma 0}\right)^2} \tag{S6}$$

where $\Delta a_{\gamma 0}$ was determined by calculating three independent $1\mu$s trajectories of single IRE1$^{516\text{-}571}$ monomer in 50% DOPC-50%POPC lipid at bilayer tension zero. $\frac{\partial \Delta E_{p1-2}}{\partial a_{\gamma 0}}$ were determined as follows

$$\frac{\partial \Delta E_{p1}}{\partial a_{\gamma 0}} = \frac{K_t}{2}\int\int\left[\frac{-2h(x,y)}{{a_{\gamma 0}}^3}\left\{h(x,y)-a_{\gamma 0}\right\}\right] dx\, dy \tag{S7}$$

$$\frac{\partial \Delta E_{p2}}{\partial a_{\gamma 0}} = \frac{K_t}{2}\int\int\left[\frac{-2a}{{a_{\gamma 0}}^3}\left\{a-a_{\gamma 0}\right\}\right] dx\, dy \tag{S8}$$

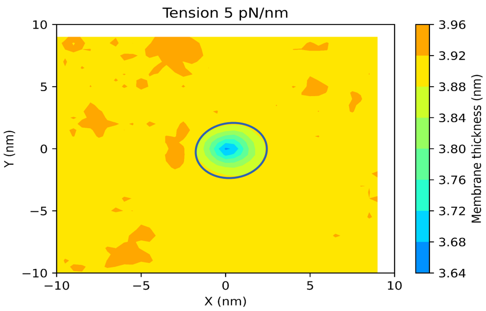


(a) Membrane thickness plot averaged over $1\mu$s simulation of single IRE1$^{516\text{-}571}$ in 50%DOPC-50%POPC under application of bilayer tension 5 pN/nm. IRE1$^{516\text{-}571}$ locally depressed the membrane. A blue curve encloses the membrane depressed by IRE1$^{516\text{-}571}$. Unperturbed membrane thickness, a was calculated by averaging the membrane area excluding the area depressed by IRE1$^{516\text{-}571}$.

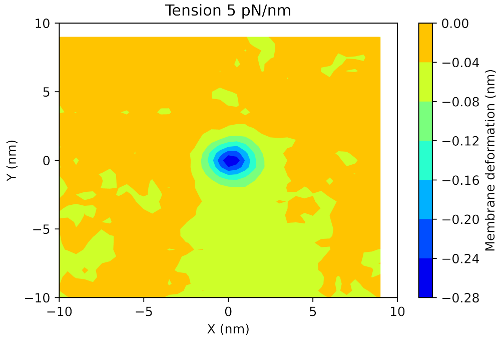


(b) Membrane deformation field of single IRE1$^{516\text{-}571}$ monomer in 50%DOPC-50%POPC averaged over 1 $\mu$s simulation at bilayer tension -5 pN/nm.

Supplementary Figure S7: Membrane thickness and membrane deformation field under application of 5 pN/nm. These membrane thickness and deformation was used to calculate $k_{\mathrm{t}}$ and $\Delta E_{\mathrm{p}}$ by using equation S3 and S4 respectively.

##### 3.3 Calculation of $\Delta E_{\gamma}$

$\Delta E_{\gamma}$ is the amount of energy required to increase the lipid area surrounding the IRE1$^{516\text{-}571}$ to conserve the ER volume. $\Delta E_{\gamma}$ can be expressed in terms of membrane deformation $h(x,y)-a$ as follows

$$\Delta E_{\gamma} = \gamma \int\int \frac{h(x,y)-a}{a_{\gamma 0}}\, dx\, dy, \tag{S9}$$

$a_{\gamma 0}$ and, $a$ were calculated in similar procedure described in section 3.2. The uncertainty of the $\Delta E_{\gamma}$ is determined as follows

$$\Delta\Delta E_{\gamma} = \left(\frac{\partial \Delta E_{\gamma}}{\partial a_{\gamma 0}} \Delta a_{\gamma 0}\right) \tag{S10}$$

where $\frac{\partial \Delta E_\gamma}{\partial a_{\gamma 0}}$ is calculated as

$$\frac{\partial \Delta E_\gamma}{\partial a_{\gamma 0}} = -\int\int \frac{\gamma}{{a_{\gamma 0}}^2} \left[h(x,y) - a\right] \, dx \, dy \tag{S11}$$

### References


A.-F. Bitbol, D. Constantin, and J.-B. Fournier. Bilayer elasticity at the nanoscale: The need for new terms. *PLOS ONE*, 7(11):1–19, Nov 2012. doi: 10.1371/journal.pone.0048306.

M. C. Watson, A. Morriss-Andrews, P. M. Welch, and F. L. H. Brown. Thermal fluctuations in shape, thickness, and molecular orientation in lipid bilayers. II. Finite surface tensions. *The Journal of Chemical Physics*, 139(8), Aug 2013. ISSN 0021-9606. doi: 10.1063/1.4818530. 084706.
